# Environmental and molecular regulation of asexual reproduction in the sea anemone Nematostella vectensis

**DOI:** 10.1101/2023.01.27.525773

**Authors:** Layla Al-Shaer, Whitney Leach, Noor Baban, Mia Yagodich, Mathew C. Gibson, Michael J. Layden

## Abstract

Cnidarians exhibit incredible reproductive diversity, with most capable of sexual and asexual reproduction. Here we investigate factors that influence asexual reproduction in the burrowing sea anemone *Nematostella vectensis*, which can propagate asexually by transverse fission of the body column. By altering culture conditions, we demonstrate that the presence of a burrowing substrate strongly promotes transverse fission. In addition, we show that animal size does not affect fission rates, and that the plane of fission is fixed along the oral-aboral axis of the polyp. Homeobox transcription factors and components of the TGFβ, Notch, and FGF signaling pathways are differentially expressed in polyps undergoing physal pinching suggesting they are important regulators of transverse fission. Gene ontology analyses further suggest that during transverse fission the cell cycle is suppressed and that cell adhesion and patterning mechanisms are downregulated to promote separation of the body column. Finally, we demonstrate that the rate of asexual reproduction is sensitive to population density. Collectively, these experiments provide a foundation for mechanistic studies of asexual reproduction in *Nematostella*, with implications for understanding the reproductive and regenerative biology of other cnidarian species.

## Introduction

Cnidarians (*e.g*., corals, anemones, jellyfish, hydrozoans) occupy a range of niches and play large roles in many ecosystems, which arguably makes investigation of their reproductive strategies important for our understanding of ocean ecology and conservation. We are interested in understanding the mechanisms that regulate asexual reproduction in this diverse group. Asexually reproducing organisms can be obligate (only reproduce asexually) or facultative (can reproduce both sexually and asexually). Although cnidarians exhibit diverse reproductive strategies, most species are capable of facultative asexual reproduction [1]. Determining how sexual and asexual reproduction are regulated is an important first step to begin understanding the mechanisms that dictate an organism’s reproductive health and how they allocate resources between alternative reproductive strategies. While sexual reproduction has been relatively well studied in cnidarians, particularly coral spawning, factors that affect asexual reproduction are not yet well characterized in any species [2], and most studies are descriptive - focusing on how asexual reproduction occurs (*e.g*., budding, fissioning) [3–5].

Environmental factors have been found to influence asexual reproduction in non-cnidarian species including stochastic environmental conditions [6], mate availability [7], seasonality [8] and population density [7,9]. Individual attributes such as physical condition [10] and body size [11,12] have also been shown to have an influence on asexual reproduction. Within cnidarians, seasonality can affect asexual reproduction in *Aurelia* [13], population density and body size have been shown to regulate the rates of bud formation in *Hydra* [14,15], and feeding has been demonstrated to increase asexual reproduction in *Hydra*, Aurelia, and *Nematostella* [4,13,14]. Efforts to identify molecular mechanisms that regulate asexual reproduction are limited to the window during which a clone is being generated. Within Bilateria, gene expression has been characterized during asexual fissioning in the annelid *Pristina leidy* [16], and functional experiments argue TGFβ, Wnt signaling, and homeobox transcription factors regulate fissioning in planaria [12,17,18]. Within Cnidaria, FGF signaling has been implicated in extracellular-matrix degradation leading to clone detachment during asexual reproduction in the coral *Pocillopora acuta* [19], and both FGF and Notch regulate bud site position and outgrowth in *Hydra* [20,21]. Despite the current efforts, there is still a lack of data about the regulators of asexual reproduction, especially within the cnidarians, arguing additional efforts in currently studied species and additional species will facilitate uncovering broad phylogenetic patterns and individual specific regulators of asexual reproduction [2].

Here we provide an initial characterization of factors that influence asexual reproduction at both the individual and population levels in the cnidarian sea anemone *Nematostella vectensis. Nematostella* undergoes asexual reproduction by transverse fission, where physal pinching leads to the separation of the lower body column (physa) which regenerates into a clonal individual [22,23] (Fig. 1a). Additionally, *Nematostella* readily spawn in laboratory culture, making it well suited for future studies aimed at investigating condition-dependent aspects of cnidarian facultative reproduction. To the best of our knowledge, this is the first study to characterize factors that affect facultative asexual reproduction in an anthozoan cnidarian. We investigated how environmental, physical, and social conditions of *Nematostella* influenced transverse fission and identified genes differentially expressed in steady state and actively fissioning animals. Lastly, we compared gene expression in high- and low-density populations which we demonstrate to have distinct rates of asexual reproduction. We found no changes in gene expression based on population density, arguing that differential gene expression does not play a large role in regulating the rate of asexual reproduction by transverse fission.

**Figure 1:**
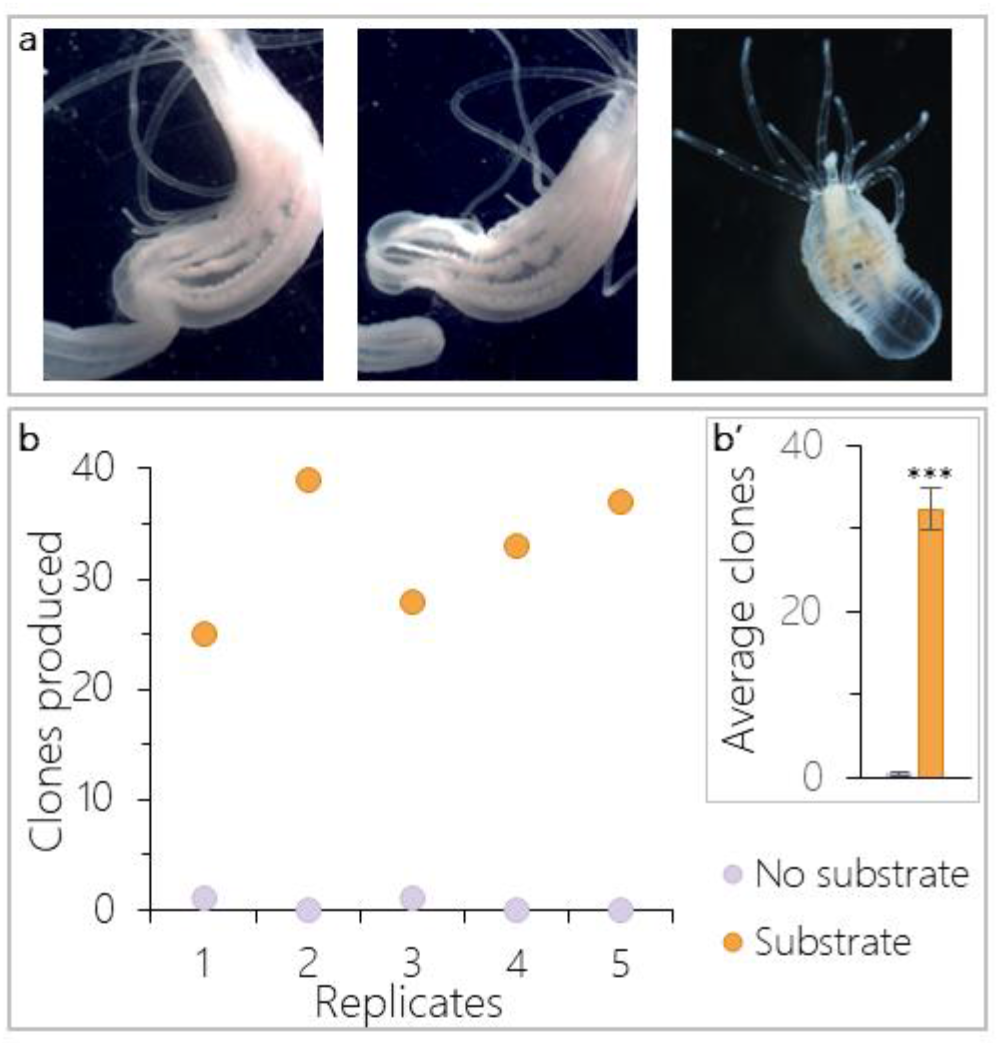
Substrate increases asexual reproduction by transverse fission. a) Transverse fission by physal pinching. Sustained constriction of the body column leads to the separation of a physal fragment which will regenerate into a functional clonal individual. b) In all replicates, more clones were produced in populations with gravel substrate than in no substrate populations and (b’) this difference was significant (t-test: *t*_(8)_ = -12.08, *p* < 0.0001, n = 5 replicates). Bars in b’ show means ± SE.

## Results

### Substrate is required for asexual reproduction in Nematostella

Asexual reproduction is rarely observed in lab populations of *Nematostella*, but it is well documented in the wild [24]. *Nematostella* is an infaunal species, but lab populations are typically maintained without substrate to burrow into, suggesting that substrate may promote transverse fissioning in *Nematostella*. To test this, we compared the number of clonal progeny produced in cultures of ten females with and without gravel substrate (n = 5 replicates). After two months, only two clonal progeny were observed within the 5 populations without substrate. In contrast, all five replicates with substrate increased by twenty five or more clonal individuals (Fig.1b, t-test: *t*_(8)_ = -12.08, *p* < 0.0001). We conclude that substrate strongly promotes fission behavior and we therefore included substrate in all subsequent asexual reproduction experiments.

To gain preliminary insight into whether there may be differences in transverse fission with and without substrate, we attempted to capture and image fissioning animals. Two time-lapse videos were created to show examples of physal pinching via transverse fission in an individual in substrate and an individual removed from substrate once the process began (Supplemental videos 1&2). The animal kept in substrate appears to rotate the body column in a twisting motion at the site of pinching until the physal fragment detaches (Supplemental video 1). The animal removed from substrate did not exhibit the same twisting motion (Supplemental video 2), indicating that the presence of substrate may provide a mechanical advantage for fissioning animals.

### Animal size does not affect fission rates

One previous study observed a highly variable latency to fission for *Nematostella* [4], but this account did not consider animal size as a potential explanatory variable. To address this possibility, a *NvLWamide-like::mCherry* neural reporter line was used to isolate animals of different sizes based on the number of longitudinal neurons each animal possessed [25,26]. Twenty nine percent of animals observed during a three-week window underwent transverse fission. Binary logistic regression showed that starting animal size did not predict whether an animal asexually reproduced (Fig. 2a; *r*^2^ = 0.005, Animal size *b* = -0.003 ± 0.008, *p* = 0.71). The latency of animals that did fission ranged from 1-18 days, and was not related to animal size (Fig. 2b, Pearson correlation: *r*_(20)_ = - 0.06, *p =* 0.80). Some individuals fissioned multiple times over the three-week period. The decrease in size after fissioning, and repeated observations of the same animal showed no consistent pattern linking animal size and latency to fission (Fig. 2b, compare points of the same color). Lastly, animals fissioned 1-4 times over a three-week period, and there was no correlation between animal size and rate of asexual reproduction (Fig. 2c; Pearson correlation: *r*_(11)_ = 0.26, *p* = 0.40). Taken together these results demonstrate that size does not determine whether an animal will fission, how fast, or how often in *Nematostella*.

**Figure 2:**
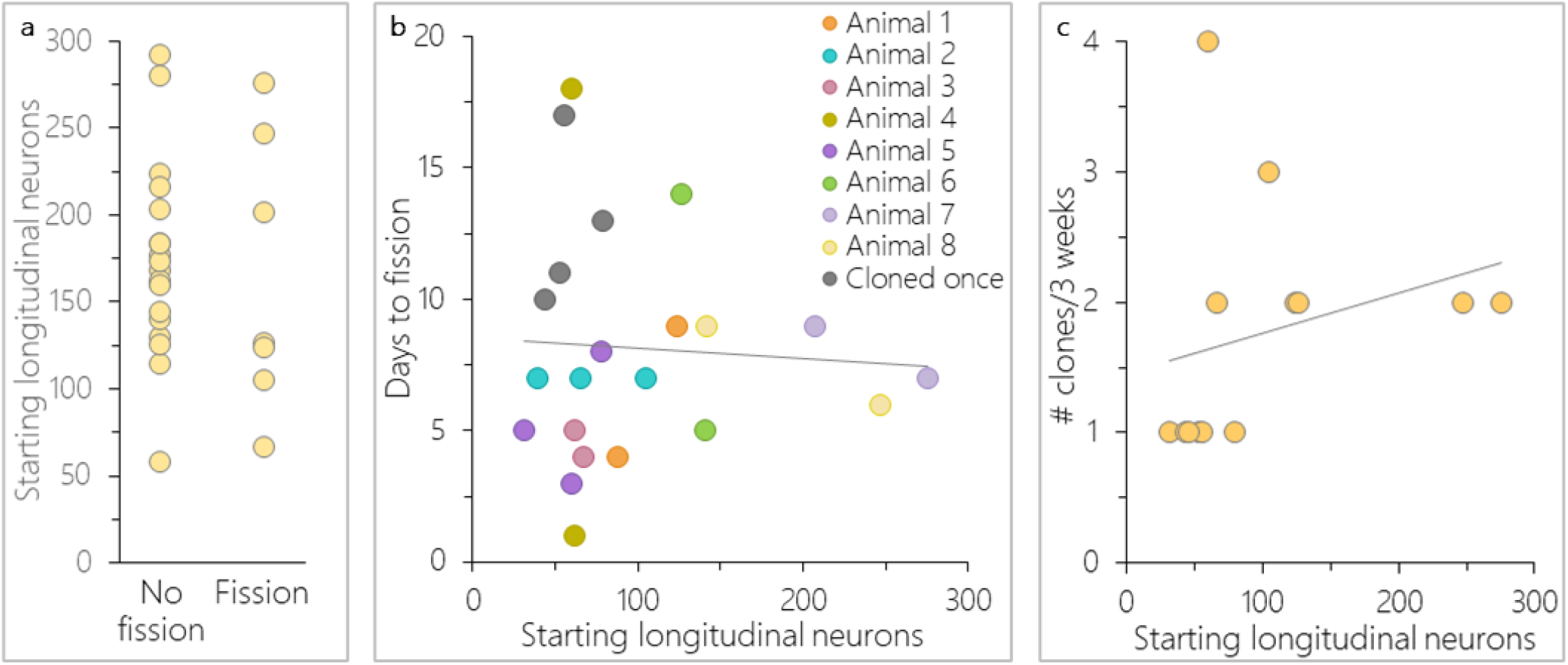
*Nematostella* size was not related to asexual reproduction. a) Only 29% of animals asexually reproduced over a three-week period. Animal size did not predict whether an animal asexually reproduced (n = 26, binary logistic regression: *r*^2^ = 0.005, Animal size *b* = -0.003 ± 0.008, *p* = 0.71). b) Latency to transverse fission varied across and within individuals, and was not related to the number of longitudinal neurons animals possessed at the start of the experiment (Pearson correlation: *r*_*(20)*_= - 0.06, *p =* 0.80; n = 12 animals and 22 fission events). c) Animal size also did not predict the rate of asexual reproduction (Pearson correlation: *r*_(11)_ = 0.26, *p* = 0.40; n = 13). Linear trendlines are shown in b&c.

### Physal pinching occurs at a set point along the body column

To determine if the physal pinching that precedes transverse fission occurs in a fixed location along the oral-aboral axis, we exploited the even distribution along the oral-aboral axis of longitudinal neurons expressing the *NvLWamide-like::mcherry* transgene [27]. The even distribution of neurons allowed us to quantify the number of neurons present within the oral (parent) and aboral (clone) remnants within < 24 hours of a fission event to determine the pre-fission size of the parent animals, and to express the position of the fission site as a percent of the total body size. The difference between parent size before and after transverse fission increased with increasing adult size, indicating that clone size scaled with parent size (Fig. 3a). However, regardless of starting size, individuals lost 26.7% of their longitudinal neurons to their clones and there was relatively low variability between individuals in this value (± 1.3% SEM) (Fig. 3b). Individuals that underwent consecutive fissioning events showed the same degree of consistency in terms of the position of the fission event along the oral-aboral axis (Fig. 3a, *e.g*., compare aqua lines from animal 2). These data suggest that the position of physal pinching during transverse fission is fixed along the oral-aboral axis of the polyp.

**Figure 3:**
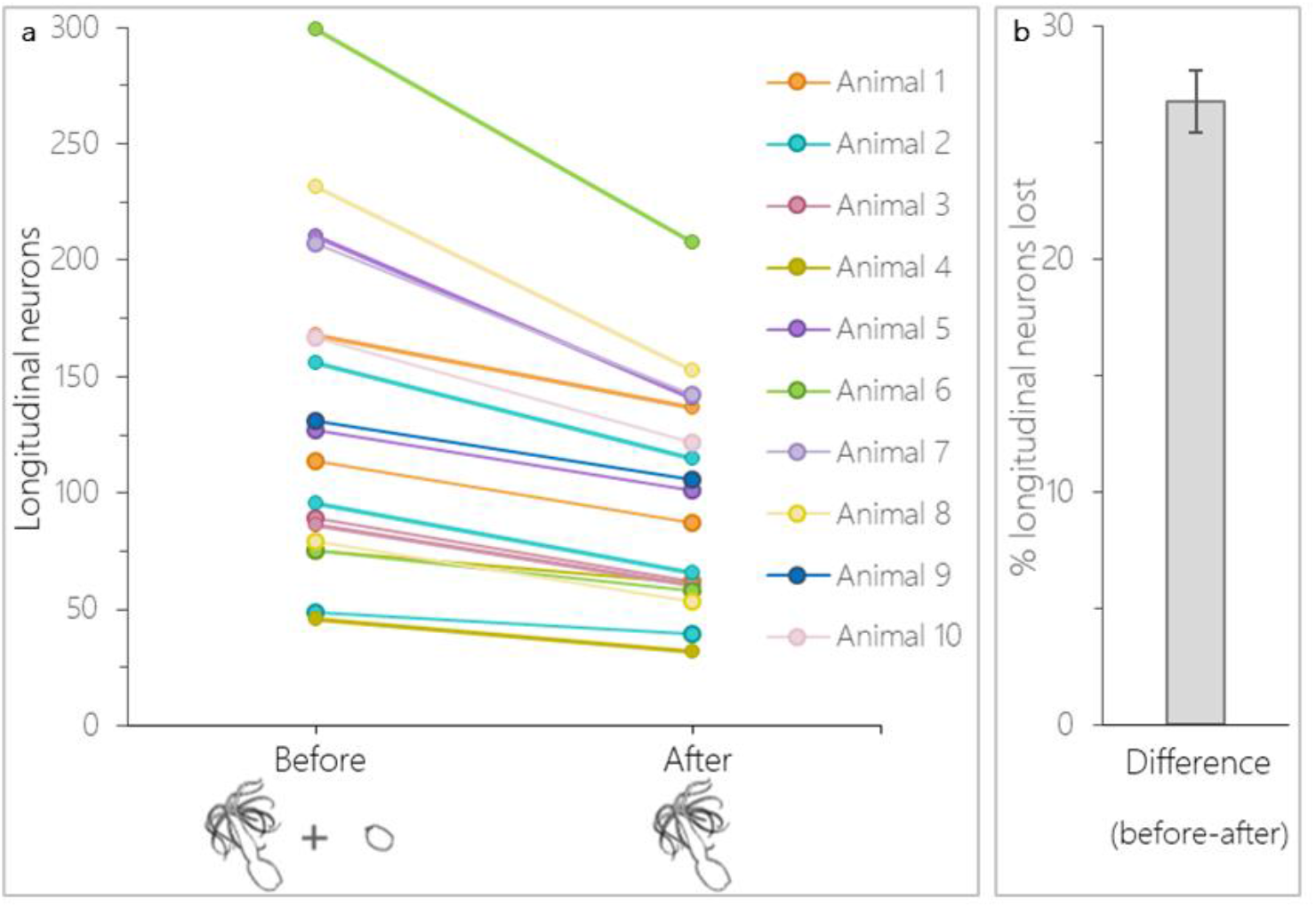
Physal pinching occurs at a set point along the body column. a) Number of longitudinal neurons before (parent + clone fragments) and after (parent only) a fission event. b) Adults decreased by an average of 26.7 ± 1.3% SE in longitudinal nerve-net size after a fission event (n = 10 animals and 18 fission events).

### Differential gene expression provides insight about mechanisms of transverse fission

Bulk RNA sequencing on fissioning (pinching) and steady state (not pinching) animals was used to identify genes differentially expressed during transverse fission. An average of 31 million reads were generated per sample which were mapped to the NVEC200 *Nematostella* genome [28]. A principal component analysis (PCA) revealed that samples segregated by pinching status along PC1, which explained 53% of the variance (Fig. 4a). A total of 1,577 differentially expressed genes (DEGs) were identified with 517 genes upregulated and 1060 downregulated in pinching animals (Fig. 4b).

**Figure 4:**
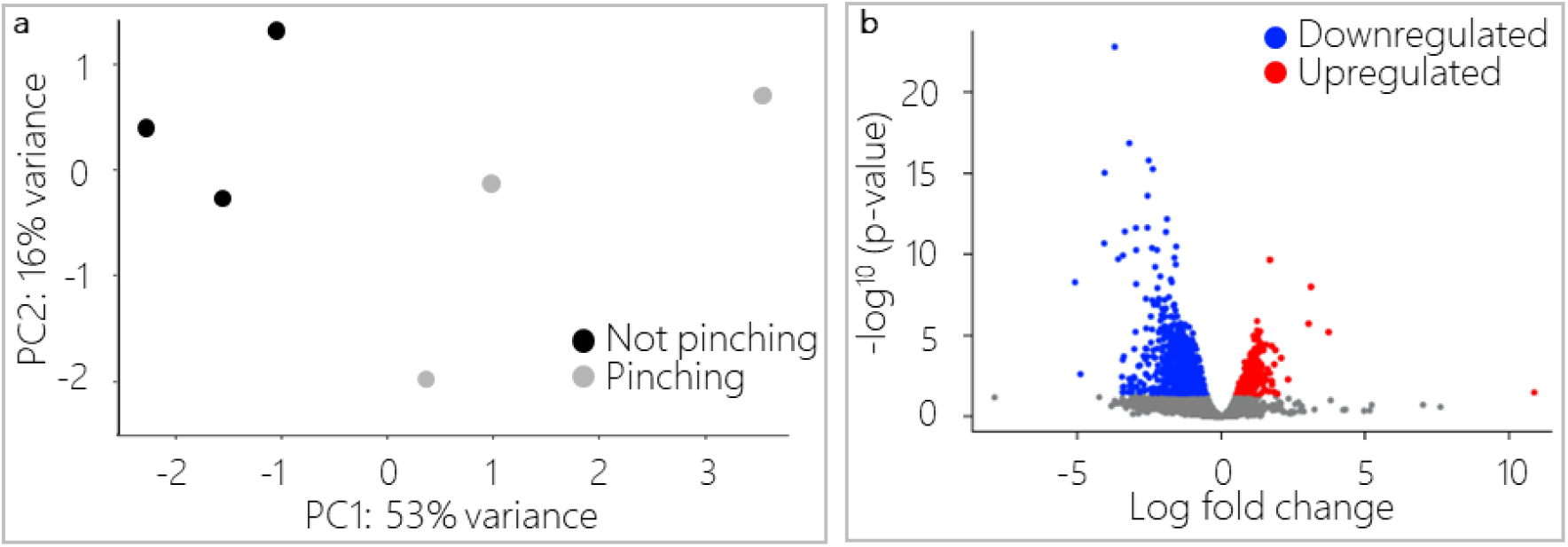
Gene expression varied between animals undergoing transverse fission and steady-state individuals. a) PCA showed that individual replicates separated along PC1 according to their pinching status. b) A total of 517 genes were upregulated and 1060 were down regulated in pinching versus steady state animals.

Within the list of DEGs we identified some transcripts known to regulate fissioning or asexual reproduction in other species (see Introduction). For example, hox genes have been implicated in transverse fission in planarians and we identified five predicted homeobox transcripts that were upregulated in pinching animals. Upregulation of TGFβ pathway components is needed for fission initiation in planaria [12], but we identified a putative TGFβ target (TGFβ-induced protein ig-h3-like) that was downregulated in our pinching animals. Similarly, elements of the notch pathway are upregulated during *Hydra* budding [21] but the five notch-like protein transcripts we found were all downregulated. Some FGF pathway components are upregulated during asexual reproduction in other cnidarians [19,20], and we identified two FGF-like ligands that were upregulated in pinching animals. However, a FGF-like ligand and multiple FGF-like receptors were down regulated, making the role of FGF in *Nematostella* fissioning less clear. Additionally, because transverse fission leads to the detachment of the physa from the rest of the body column, we predicted that transcripts related to cell adhesion and extracellular matrix organization would be differentially expressed. As predicted, transcripts that may promote cell adhesion were down regulated, and transcripts that regulate tissue organization and cytoskeletal architecture were differentially expressed. See supplemental table 1 for transcript IDs and annotations. Together these data imply that downregulation of transcripts related to maintaining body integrity likely aid in the separation of the physal fragment and suggest roles for major developmental signaling pathways in the *Nematostella* transverse fission process.

### Gene ontology analysis uncovered functional characteristics of transverse fission behavior

To provide a broader understanding of how fissioning is regulated, a rank-based Gene Ontology (GO) enrichment analysis of the DEGs found between pinching and non-pinching animals was conducted. Using a cutoff of *p ≤* 0.05, we identified 23 cellular component terms (17 up, 6 down), 49 molecular function terms (32 up, 17 down), and 60 biological process terms (45 up, 15 down) (Supplemental figures 1-3). The top upregulated biological processes (*p* ≤ 0.001) included negative regulation of cell cycle, DNA replication, and RNA metabolism (Fig. 5). The top downregulated biological process terms included cell adhesion, developmental processes, and intracellular signal transduction (Fig. 5). Collectively these findings suggest that during fissioning the cell cycle becomes tightly regulated, most likely suppressed, while cell adhesion and patterning mechanisms are downregulated to promote effective separation of the physa from the body column. Additional studies are needed to fully understand how the genes identified in this study regulate transverse fissioning.

**Figure 5:**
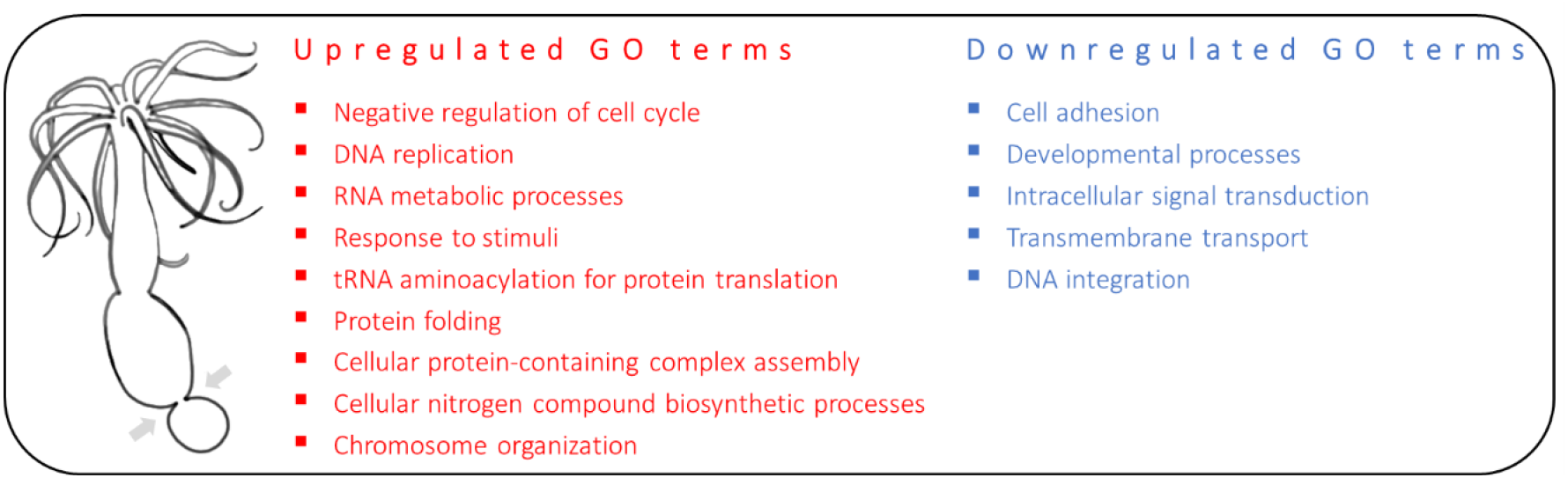
Biological processes implicated during physal pinching. Rank-based GO analysis of top (*p* ≤ 0.001) upregulated and downregulated biological processes during physal pinching. Based on DEGs identified between pinching and non-pinching treatments.

### Asexual reproduction increases at low population density

An overarching hypothesis in species with facultative reproduction is that there should exist specific environmental conditions that act to promote asexual reproduction, otherwise it is difficult to conceive of how multiple modes of reproduction could be evolutionarily maintained [6,29]. In the wild, tidal events can collect and trap *Nematostella* in isolated salt marsh pools at varying population densities. We hypothesized that population density would affect rates of asexual reproduction and predicted that asexual reproduction would increase in animals kept at low-density due to reduced mating opportunity. Animals were placed into either a high or low-density population (40 or 4 animals respectively). After three weeks, the number of clones produced per starting individuals was determined for each density treatment. Although there was variation in the overall number of clones produced, with the least asexual reproduction occurring April-May, the effect of density was always the same for all replicates. Animals at low-density asexually reproduced more than those in high-density treatments (Fig. 6a). On average, individuals from low-density populations generated significantly more clones than those from high-density populations (Fig. 6a’; t-test: *t*_(32)_ = 3.2, *p* = 0.003).

**Figure 6:**
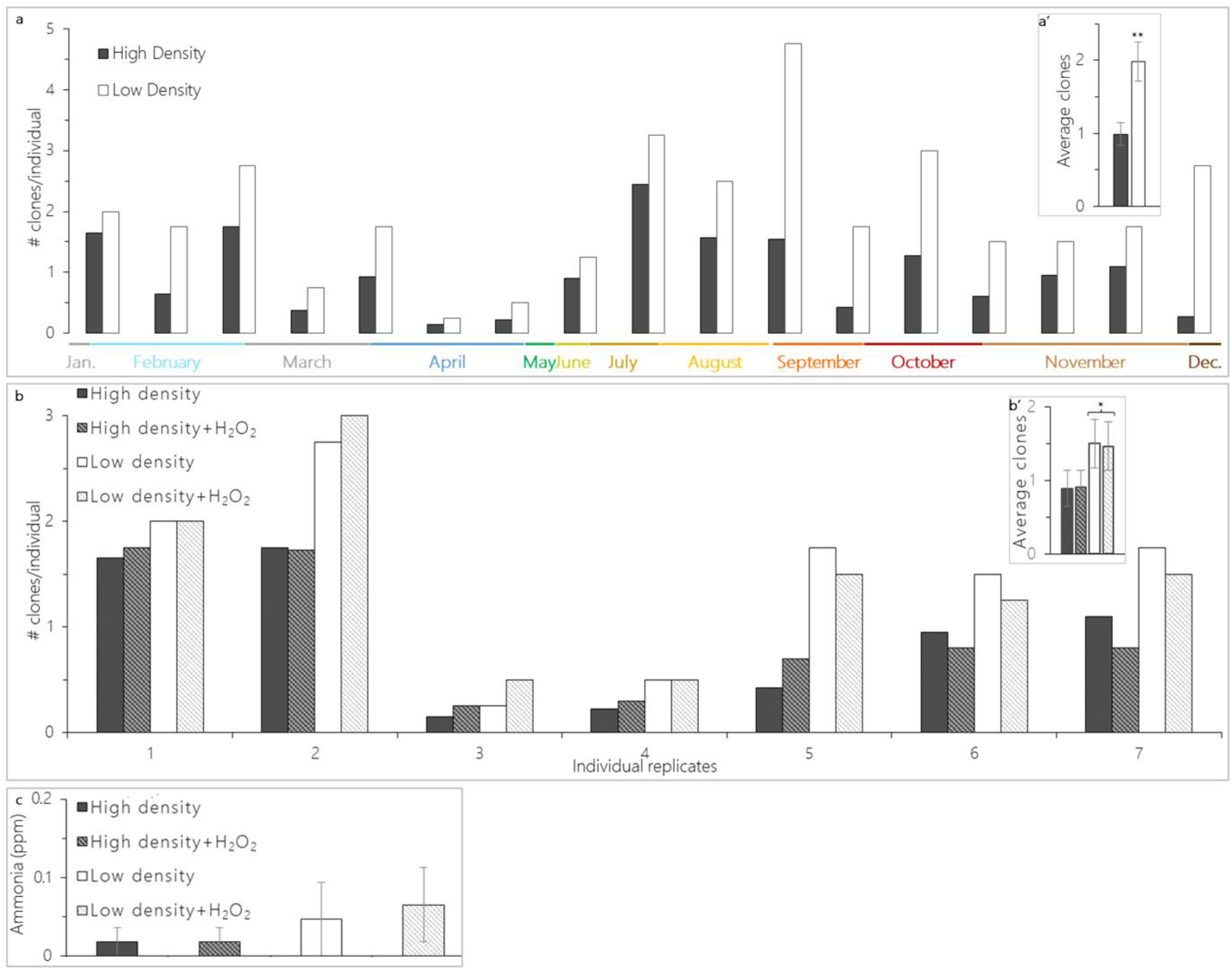
Population density affects *Nematostella* asexual reproduction. a) High- and low-density animal populations (40 or 4 animals respectively) were kept in large petri dishes (a’) during the experiment period. b) Regardless of time of year, all replicates generated more clones per starting individual in low-versus high density treatments. b’) On average there was a significant difference in asexual reproduction between density treatments (t-test: *t*_(32)_ = 3.2, *p* = 0.003; n = 17 replicates). c) Regardless of H_2_O_2_ presence, nearly all replicates had more clones generated in both low-density treatments compared to both high-density treatments. c’) On average, there was a significant difference in asexual reproduction based on density and this was not influenced by H_2_O_2_ treatment (mixed ANOVA – main effect density: *F*_(1,24)_ = 4.1, *p* = 0.05; H_2_O_2_ X density: *F*_(1,24)_ = 0.007, *p* = 0.94; n = 7 replicates). d) Ammonia levels did not differ between treatments (mixed ANOVA-H_2_O_2_ X density: *F*_(1,24)_ = 0.06, *p* = 0.81; n = 7 replicates). Bars in b’, c’, and d show means ± SE.

We questioned if differences in metabolic waste buildup might explain the density-based differences in asexual reproduction. To test this, we repeated the above experiment but added additional low- and high-density treatments that received H2O2 in their water (final concentration 0.00025% or 82µM). This concentration is not lethal and does not inhibit regeneration in *Nematostella* [30]. Like before there was variability in the amount of asexual reproduction that occurred, but in nearly all replicates low-density animals produced more clones than high-density, regardless of whether H_2_O_2_ was present (Fig. 6b). On average, there was an overall effect of density. Both low-density treatments produced significantly more clones than both high-density treatments (Fig. 6b’; mixed ANOVA – main effect density: *F*_(1,24)_ = 4.1, *p* = 0.05) and this was not influenced by the addition of H_2_O_2_ (mixed ANOVA - H_2_O_2_ X density: *F*_(1,24)_ = 0.007, *p* = 0.94). The amount of ammonia present across all treatments was also measured to determine if differences in this nitrogenous waste might explain density-based differences. Ammonia levels were low and did not differ between any treatments (Fig. 6c; mixed ANOVA-H_2_O_2_ X density: *F*_(1,24)_ = 0.06, *p* = 0.81). We conclude that *Nematostella* can sense their population density and then adjust their asexual reproduction accordingly. The mechanism by which this occurs is unknown, but differences in the metabolic waste products ammonia and H2O2 are not responsible.

### Gene expression did not vary based on population density

To better understand the effect that density has on *Nematostella*, we performed bulk mRNA sequencing on whole individuals collected from low- and high-density treatments at the end of the experiment period. None of the sampled animals were undergoing physal pinching at the time of collection. We predicted that there would be differential gene expression based on density treatment alone, but that differentially expressed genes might not be related to the process of asexual reproduction since this process is unpredictable and asynchronous across individuals. Sequencing yielded an average of 50 million raw reads per sample which were mapped to the *Nematostella* NVE transcriptome v2.0 [31]. Using transcripts per million normalized read counts, the average distance between samples was estimated using a PCA that showed that samples did not separate along PC1 (60% variance) or PC2 (11% variance) based on population density treatment (Fig. 7a). Further, a differential gene expression analysis showed that no transcripts were differentially expressed between samples from high- and low-density treatments (Fig. 7b), indicating that within our experimental context changes in gene expression does not explain density-based differences in asexual reproduction.

**Figure 7:**
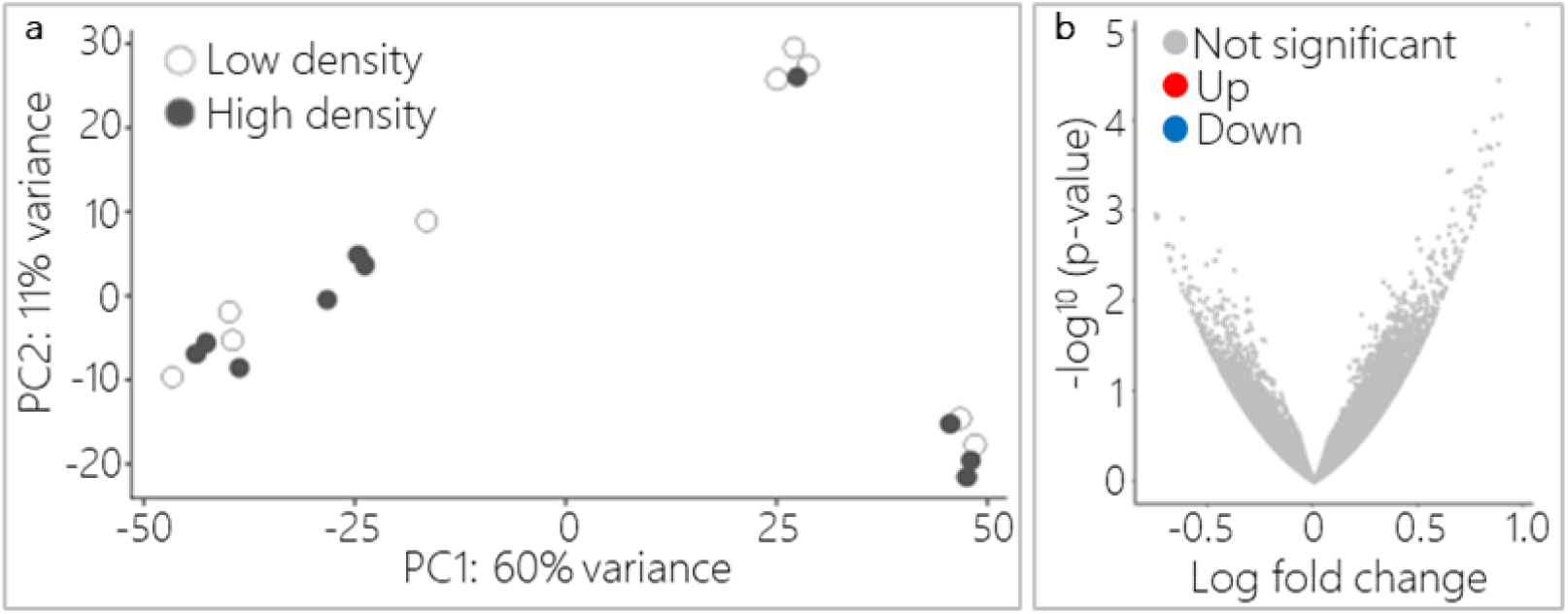
Gene expression did not differ based on population density treatment. a) PCA revealed that replicates did not segregate by density treatment along PC1 or PC2 (low-density n = 9, high-density n = 10). b) DeSeq2 analysis showed that no transcripts were significantly different in expression between low- and high-density samples (all adjusted *p* > 0.05).

## Discussion

One of our primary goals was to expand our knowledge of cnidarian reproduction by characterizing affecters of asexual reproduction in the cnidarian *Nematostella*. It was not surprising that substrate enhanced and is effectively required for transverse fission to occur since they are naturally burrowing anemones. Our preliminary observations that animals that fission in substrate appear to twist, make it enticing to speculate that substrate offers a mechanical advantage to promote fissioning success. However, it is not clear if the attempts at fissioning are similar in substrate compared to no substrate cultures and substrate could also play a larger role by increasing the rate of physal pinching that leads to transverse fission. Regarding animal size, we did not detect any size requirement related to transverse fission behavior. Since when tracking individual animals there was variability in whether transverse fission occurred, there must be other factors that influence whether or not animals in isolation fission. We speculate that body condition, primarily due to food intake prior to the start of the experiment, is a likely factor because increases and decreases in feeding are known to influence fission rates in *Nematostella* [4].

The rate of *Nematostella* asexual reproduction by transverse fission increases in populations at lower density. This observation argues that animals are capable of sensing the number of other anemones and increase their rates of asexual reproduction to increase fitness when competition from other animals is low or mating potential is decreased. This finding presents a number of obvious questions. For example, how do *Nematostella* sense population density and subsequently alter fission rates, or do sex ratios influence asexual reproduction rates at different population densities? Some insights come from our observations. There was no difference in gene expression between animals at high- and low-density which argues that density does not alter transcription rates, suggesting an alternative mechanism for density dependent asexual reproduction rates. One possibility is that *Nematostella* use sensory information to interpret their social environment and in turn regulate their fission behavior. *Nematostella* lack vision, and it is unlikely that density is determined by touch since burrowed adults rarely move around and individuals were placed equidistant from one another in our experiments. Our favored hypothesis is that detection of a chemical signal in the environment initiates a cascade of events that mediate physal pinching and subsequently transverse fission. This hypothesis is supported by the fact that chemical communication is a common sensory modality in aquatic organisms [32,33] and it has been implicated in the initiation and synchronization of spawning in corals [34–37]. The chemical signal is likely not differences in metabolic waste buildup since there was no effect of H2O2 or ammonia on density-based differences in fission rates, but we acknowledge that other waste products may be co-opted for discerning population density.

Regardless of what regulates asexual reproduction, we know that it must have a downstream effect on initiating physal pinching – which we have identified to occur at a set point along the oral-aboral axis (∼ bottom 27% of parent is lost to clone). Set fission locations have also been identified in bilateria [12] and in *Hydra*, where budding occurs in the bottom 1/3^rd^ of the body column [38]. This similarity between *Hydra* budding location and *Nematostella* physal pinching location is intriguing because it suggests that there may be a conservation of mechanisms that designate the asexual field in cnidaria. It is unknown what causes physal pinching at this set location. In planaria, the pinching that precedes fissioning is neurally regulated [11,12], and it is enticing to speculate that in *Nematostella* neural regulation controls localized constriction by causing columnar ring muscle to constrict [39] at the pinching site.

Another goal was to establish a foundational framework for further exploration of the molecular mechanisms of transverse fission behavior. Once *Nematostella* are actively fissioning, changes in gene expression occur which presumably lead to the detachment of the physa from the body column. We identified 1577 transcripts that were differentially expressed between pinching and steady state animals, including predicted homeobox and notch-like protein transcripts as well as TGFβ and FGF pathway components – all of which have been implicated in asexual reproduction in other bilaterians and cnidarians (see Introduction). Functional experiments and further characterization of differential gene expression over time throughout the fission process are needed to improve the molecular model of physal pinching in *Nematostella*, and to determine if there is conservation between taxa. For example, FGF and notch signaling help define the boundary between bud and parent in *Hydra* [20,21] and it is possible that these pathways could have a similar role in *Nematostella*. Additionally, DEGs related to the biological processes of negative cell cycle regulation and cell adhesion should be further explored. The downregulation of the latter probably aids in detachment of the physa, especially given that FGF signaling has been linked to extracellular-matrix degradation and subsequent asexual polyp detachment in the coral *Pocillopora acuta* [19].

Our findings lend insight into whether the factors and mechanisms that govern asexual reproduction are conserved across the Cnidaria but continued work in *Nematostella* will allow us to draw broader conclusions. The potential to use *Nematostella* as a model to identify the conditions that promote asexual versus sexual reproduction to better our understanding of cnidarian facultative reproduction should also be explored. It is our opinion that any improvement on our knowledge of the proximate and ultimate factors that govern cnidarian reproduction are of value because it will allow us to better understand a phylum that exhibits incredible reproductive diversity, plays essential ecological roles, and can be applied to management plans or research into strategies aimed to bolster declining wild populations.

## Methods

### Animal care and relaxation paralytic

*Nematostella* were maintained in mixed sex groups kept in glass bowls filled with1/3X artificial salt water (1/3X ASW; pH 8.1-8.2) and kept in a dark 17^°^ C incubator. Freshly hatched artemia were fed four times a week and water changes occurred weekly. Since it is known that asexual reproduction is suppressed when *Nematostella* are food deprived [4], enough artemia was fed to prevent degrowth [27] and was controlled within each experiment. When necessary, animals were relaxed with MgCl_2_ (7.14% Wt/Vol) in 1/3X ASW before length measurements or neuron counts were taken.

### Substrate and asexual reproduction

For this experiment a colony of adult female *Nematostella* were clonally propagated in glass dishes over several generations in the laboratory as described by Hand and Uhlinger [40] and were formally naive to substrata. One hundred individuals of similar body size were subset evenly into ten groups and placed in two-cup Pyrex dishes containing either gravel substrate (n = 5) or no substrate (n = 5). The dishes and gravel were washed and autoclaved prior to usage. After placement of animals into their respective conditions, the experiment was left to run continuously for two months. During this time, animals were fed green mussel three times weekly followed by weekly water changes where old water was decanted and replaced with fresh 1/3X ASW. The substrate was never removed or washed during the experiment. At the conclusion of the study, animals were re-counted using a mesh window screen to sift through the substrate. Videos of fission behavior were generated from time lapse images taken at five-minute intervals of an individual kept in substrate and an individual removed from substrate, after pinching had begun, over the course of eleven and four hours respectively. Imaging was done with either an sCMOS camera with an attached lens (Thor Labs Quantalux CS2100M-USB and MVL5M23 respectively) or a Nikon dissection microscope (Nikon SMZ1270) with a USB camera attachment (Levenhuk M300). To help visualize fission behavior of the individual kept in substrate, a clear polydimethylsiloxane (PDMS) based substrate was made. The PDMS (Sylgard184 Ellsworth Adhesives 184 SIL ELAST KIT 0.5KG) was mixed at a ratio of 10:1 (prepolymer to curing agent), degassed until clear, and cured at 80° C for twenty minutes before cutting the solid slab into small pieces.

### Animal size and asexual reproduction

Longitudinal neurons were counted in *NvLWamide-like::mCherry* expressing animals at the start of the three-week long experiment period for the latency to fission, rate of fission, and fission location experiments. Neurons were counted by hand on live animals relaxed with MgCl_2_ using a dissection scope equipped with fluorescence (Nikon SMZ 1270) [27]. After counting, *Nematostella* were individually placed in petri dishes (100 × 15mm) filled with 38g gravel substrate and 38 mL 1/3X ASW and kept in a 22^°^ C incubator with a 12:12 light-dark cycle. Replicates were fed 10% of their starting longitudinal neuron count in artemia twice a week, and each replicate received a water change weekly. Individuals were checked daily for clones by gently looking through the substrate under a dissection scope. If asexual reproduction had occurred, the adult and clone fragments were relaxed with MgCl_2_ and longitudinal neurons were counted.

### Transcriptomic analysis of animals during physal pinching

Animals were placed in gravel substrate and individuals were either sampled as physal pinching occurred (pinching) or in steady state animals (not pinching) (n = 3 pooled anemones per treatment). Animals were put into RNAlater and stored at -20°C until RNA isolation. Total RNA was isolated from all samples using the RNAqueous kit (Ambion) followed by DNAse treatment with a DNA-free kit (Invitrogen). RNA quantity was assessed using both a Qubit Fluorometer with associated RNA Broad Range kit (Thermo Fisher Scientific) and an Agilent 2100 Bioanalyzer using a “pico” kit, each according to their respective manufacturer’s instructions. Libraries were prepared from 14.6-100ng of high-quality total RNA by the Sequencing and Discovery Genomics core at the Stowers Institute for Medical Research following manufacturer protocols for the Illumina TruSeq Stranded mRNA Library Prep kit and Illumina TruSeq RNA Single Indexes Sets A and B. Resulting short fragment libraries were checked for quality and quantity using an Agilent Bioanalyzer and a Qubit Fluorometer. Libraries were pooled at equal molar concentrations, quantified, and sequenced as 75bp single reads on a high-output flow cell using the Illumina NextSeq 500 instrument and NextSeq Control Software 2.2.0.4. Following sequencing, Illumina Primary Analysis version RTA 2.4.11 and bcl2fastq2 v2.20 were run to demultiplex reads for all libraries and generate FASTQ files. Sequencing produced an average of 31,062,832 million reads per sample, ranging from 26,312,701 to 33,855,476. Sequencing results were deposited in the NCBI GEO database (GSE223794). The filtered reads were aligned to the Nvec200 *Nematostella* genome (Zimmermann et al. 2022) with STAR aligner (version 2.7.3a) using Ensembl 99 gene models. Reads mapping to the reference at the same start position with 100% alignment identity of the transcript were regarded as PCR duplicates and discarded. TPM values were generated using RSEM (version v1.3.0).

All statistical analyses were completed in the R environment (R3.5.0). The raw read-counts-per-gene table was filtered to contain only genes with at least three counts and used to perform outlier detection with the arrayQualityMetrics R package [41] with three different outlier detection methods: *distance between arrays, boxplots*, and *MA plots*. No samples were excluded from downstream analysis. The filtered count data were analyzed using the R package DESeq2 [42], which produced a normalized counts-per-gene table used in subsequent analyses. Normalized count data were transformed using the *rlog* function. Corrections were made for multiple testing at a cutoff of 0.1. Using a generalized linear model with Wald pairwise comparisons, DEGs were determined between samples that were pinching or not pinching. The VEGAN package was used to visualize and calculate significance and generate a PCA of variance-stabilized gene expression [43]. Functional summaries of differentially expressed genes were determined by rank-based Gene Ontology (GO) enrichment analysis, using signed, unadjusted log-transformed p-values (positive if up-regulated, negative if down-regulated) with the GO_MWU package (https://github.com/z0on/GO_MWU). This method uses a Mann-Whitney U test and measures whether each GO term is significantly enriched in up- or down-regulated genes based on their delta rank (quantitative shift in rank) rather than looking for GO terms among significant genes only.

### Population density and asexual reproduction

Lab cultured wild type *Nematostella* from mixed sex populations were used for all population density experiments. Prior to being assigned to a high or low-density treatment (n = 40 or 4 animals respectively), all animals were relaxed in MgCl_2_ and their body length was measured using a clear plastic ruler placed under their petri dish. Within each replicate, average animal length and length range was controlled across population density treatments. Animals were placed approximately equidistant in their assigned treatment petri dish (145 × 25mm) filled with 100g gravel substrate and 200ml 1/3X ASW and kept in a 22^°^ C incubator with a 12:12 light-dark cycle for the three-week long experiment period. High- and low-density treatments were fed 400 and 40 µl artemia respectively, twice a week. Water changes were performed weekly and a 20mL water top off occurred mid-week. H_2_O_2_ density treatments had H_2_O_2_ added to their 1/3X ASW every time water was added (final concentration 0.00025% or 82µM). Replicates were checked daily for egg clutches which were removed if found. Ammonia levels were tested on water removed from each treatment dish during weekly water changes using an API brand NH_3_/NH_4_ ^+^ test kit. At the end of the experiment period, all animals were removed from each dish and counted to determine the number of clones that were generated per starting individual. Dishes were checked multiple times under a dissection scope to ensure no animals were missed.

### Transcriptomic analysis of population density animals

Total RNA was extracted from animals at the end of the population density experiment period. In total ten high- and nine low-density animals were collected from four different population density replicates. None of the sampled individuals were undergoing physal pinching at the time of collection. Each animal was flash frozen in liquid nitrogen, processed with a liquid-nitrogen cooled mortar and pestle, stabilized in 1mL TRI Reagent (Sigma-Aldrich), and then stored at -80^°^ C until. Total RNA was isolated as previously published [44] RNA concentration and purity were measured using a NanoDrop spectrophotometer (all samples OD:260/280 and OD:260/230 ≥ 2) and sample integrity was confirmed using an Agilent 2100 Bioanalyzer (all samples RIN ≥ 8.0) following manufacturer instructions. RNA was stored at -80^°^ C until needed. Sample cDNA libraries were prepared from an average of 21-274ng of high-quality RNA. Library preparation and sequencing was done by Novogene Inc. Sample libraries were constructed following manufacturer protocols using an Illumina Stranded mRNA Prep kit, which uses poly A enrichment to purify mRNA from total RNA, and an Illumina NovaSeq 6000 was used for sequencing (paired end, 150bp reads). An average of 50,324,344 million reads were produced per sample, ranging from 38,628,894 to 70,726,914. Reads were quality filtered and adapter trimmed using Trimmomatic [45]. A read was discarded if there was adapter contamination, if it contained more than 10% uncertain nucleotides, or if there were more than 50% low quality nucleotides (Qscore ≤ 5). Sequencing data can be retrieved from the NCBI SRA database (PRJNA925453).

Sequencing data were uploaded to the Galaxy web platform, and the public server at usegalaxy.org was used to map and analyze the sequencing data [46]. Filtered reads were mapped and quantified using the Salmon quant tool with default parameters (version 1.5.1) (Patro et al. 2017) using the *Nematostella* NVE transcriptome v2.0 [31]. Transcripts per million was used to quantify transcript abundance. The count tables generated with Salmon quant were used for differential transcript analysis between high- and low-density samples using the Bioconductor DESeq2 tool using default settings (version 2.11.40.7) [42]. A false discovery rate of *p ≤ 0.05* was used to determine differential expression (adjusted for multiple testing using the Benjamini-Hochberg procedure).

## Acknowledgements

We would like to thank Jamie Havrilak and Dylan Faltine-Gonzalez for providing input and support throughout the course of this study. We are grateful to MingHe Cheng and Nolan Jetter for assisting with experiment maintenance, Jeffrey Lange for their support in generating time-lapse footage, SIMR Computation Biology Core for aiding in sequencing, and Austen Barnett for their advice regarding sequence analysis. We

