## Supplemental figures 1-3 and supplemental table 1 for "Environmental and molecular regulation of asexual reproduction in the sea anemone Nematostella vectensis"

**Supplemental video 1: Transverse fission behavior of an individual in PDMS substrate**. Time lapse images were captured every 5 minutes for 11 hours, video speed is ~ 4 FPS.

**Supplemental video 2: Transverse fission behavior of an individual removed from substrate after pinching began.** Time lapse images were captured every 5 minutes for 4 hours, video speed is ~ 4 FPS.

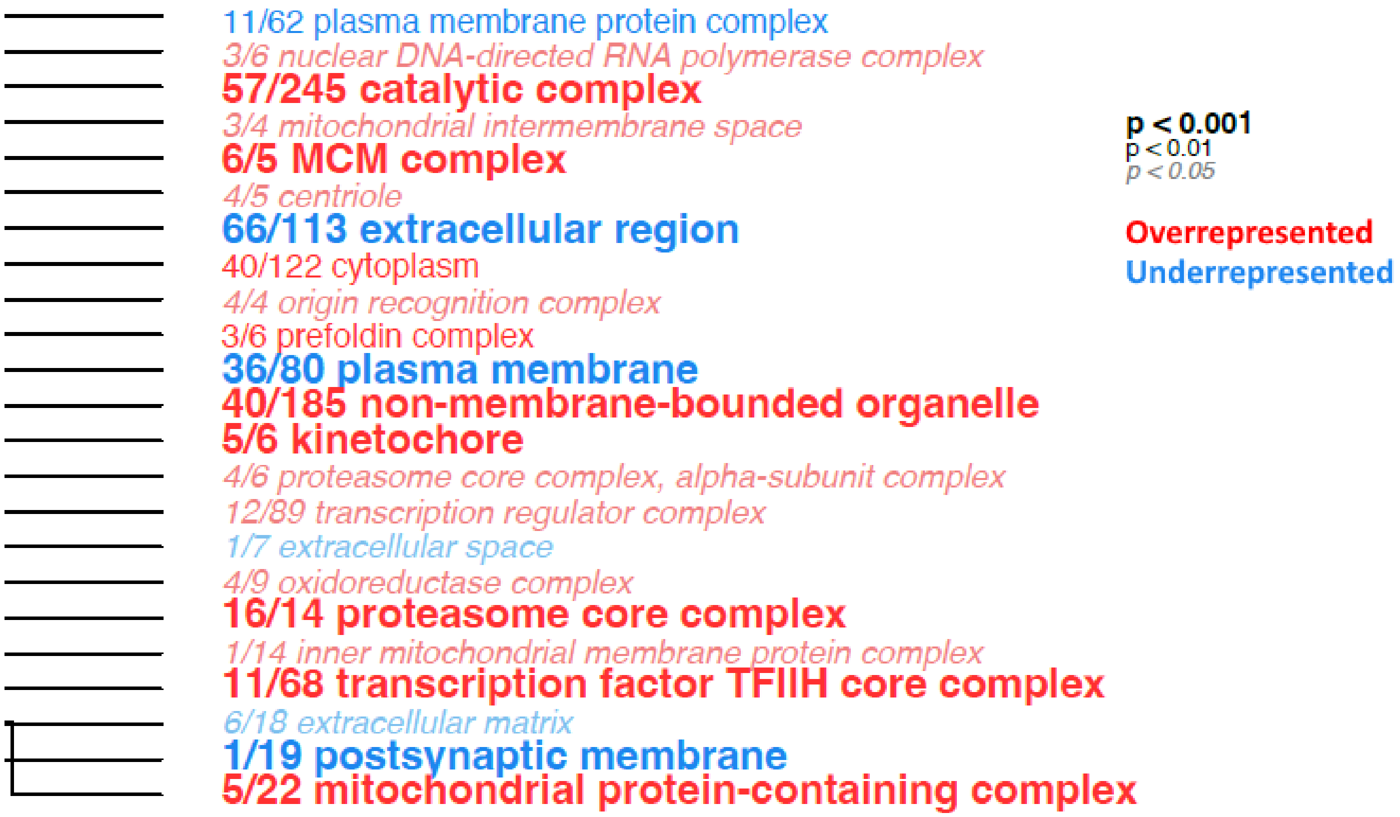


**Supplemental figure 1: Rank-based GO enrichment analysis of cellular components in pinching versus steady state animals.** Fractions indicate the Number of DEGs matching the GO term/Total number of genes matching the GO term. Overrepresented and underrepresented terms are colored red and blue respectively and significance is indicated by text size.

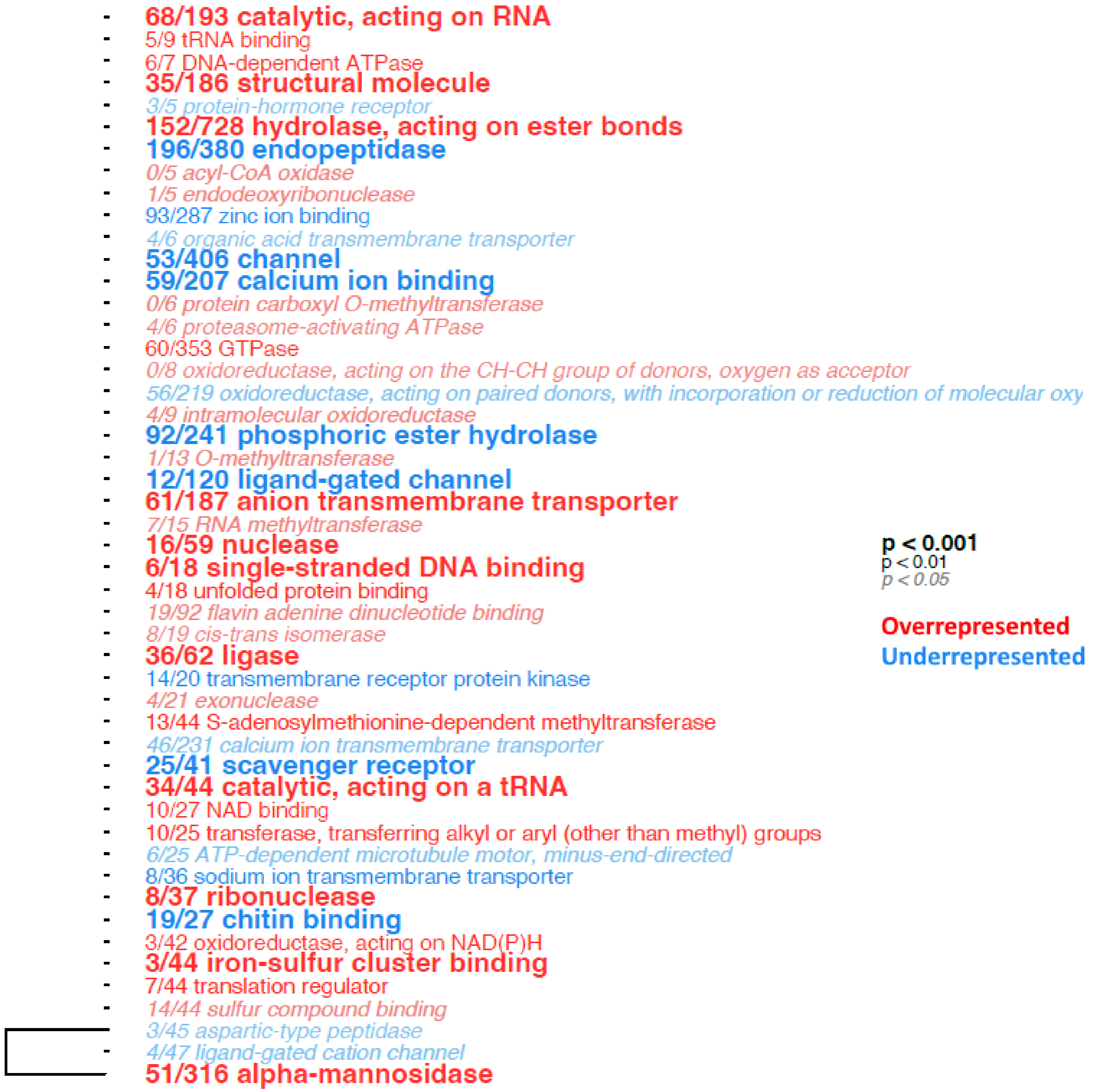


**Supplemental figure 2: Rank-based GO enrichment analysis of molecular functions in pinching versus steady state animals.** Fractions indicate the Number of DEGs matching the GO term/Total number of genes matching the GO term. Overrepresented and underrepresented terms are colored red and blue respectively and significance is indicated by text size.

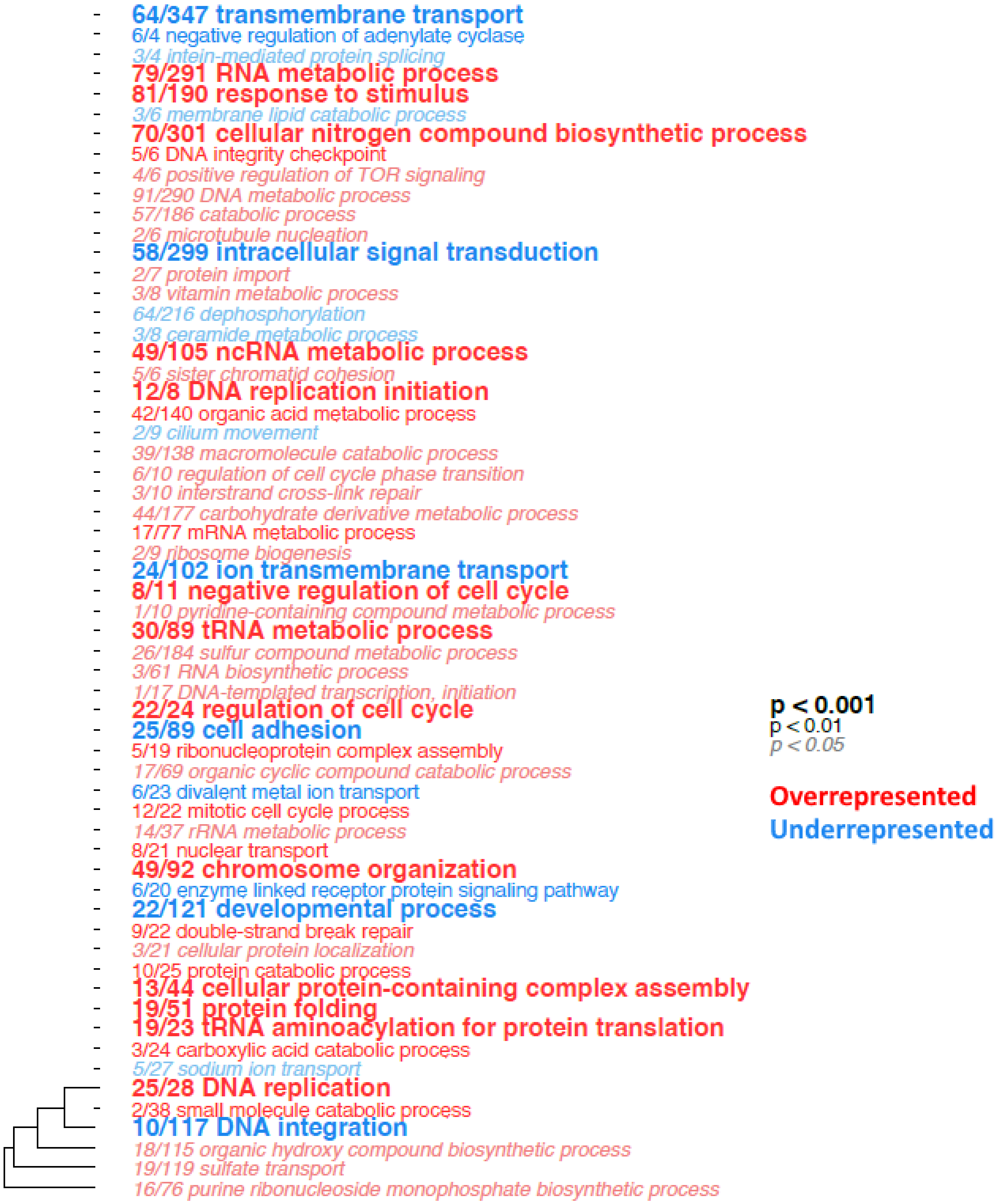


**Supplemental figure 3: Rank-based GO enrichment analysis of biological processes in pinching versus steady state animals.** Fractions indicate the Number of DEGs matching the GO term/Total number of genes matching the GO term. Overrepresented and underrepresented terms are colored red and blue respectively and significance is indicated by text size.

Supplemental table 1: NVEC200 transcript IDs and annotations for select differentially expressed genes in pinching versus steady state animals.

| Transcript ID | Upregulated or downregulated? | Annotation |
|---|---|---|
| NV2t011935001.1 | upregulated | T-cell leukemia homeobox protein 2 (*Mus musculus*) |
| NV2t011262001.1 | upregulated | Homeobox protein Nkx-2.2a (*Danio rerio*) |
| NV2t012726001.1 | upregulated | Homeobox protein SIX3 (*Gallus gallus*) |
| NV2t025278001.1 | upregulated | Homeobox protein B-H1 (*Drosophila melanogaster*) |
| NV2t011140001.1 | upregulated | Homeobox protein XENK-2-like (*Xenopus laevis*) |
| NV2t008483001.1 | upregulated | Fibroblast growth factor 2 (*Monodelphis domestica*) |
| NV2t006376001.1 | upregulated | Fibroblast growth factor 1 (*Notophthalmus viridescens*) |
| NV2t006385001.1 | upregulated | FRAS1-related extracellular matrix protein 2 (*Mus musculus*) |
| NV2t020441001.1 | downregulated | Transforming growth factor-beta-induced protein ig-h3 (*Mus musculus*) |
| NV2t003830001.1 | downregulated | Fibroblast growth factor receptor 3 (*Gallus gallus*) |
| NV2t019098001.1 | downregulated | Fibroblast growth factor receptor 1 (*Homo sapiens*) |
| NV2t012077001.1 | downregulated | Fibroblast growth factor receptor 3 (*Xenopus laevis*) |
| NV2t019096001.1 | downregulated | Fibroblast growth factor receptor 3 (*Gallus gallus*) |
| NV2t002934001.1 | downregulated | Fibroblast growth factor receptor 2 (*Gallus gallus*) |
| NV2t019095001.1 | downregulated | Fibroblast growth factor receptor 3 (*Gallus gallus*) |
| NV2t012704001.1 | downregulated | Fibroblast growth factor 10 (*Mus musculus*) |
| NV2t006022001.1 | downregulated | Neurogenic locus notch protein (*Drosophila melanogaster*) |
| NV2t005916001.1 | downregulated | Neurogenic locus notch protein (*Drosophila melanogaster*) |
| NV2t006444001.1 | downregulated | Neurogenic locus notch protein (*Drosophila melanogaster*) |
| NV2t019821001.1 | downregulated | Neurogenic locus notch homolog protein 1 (*Danio rerio*) |
| NV2t018428001.1 | downregulated | Neurogenic locus notch homolog protein 2 (*Mus musculus*) |
| NV2t011625001.1 | downregulated | Protocadherin-like protein (*Acropora millepora*) |
| NV2t014198001.1 | downregulated | Protocadherin Fat 3 (*Rattus norvegicus*) |
| NV2t021994001.1 | downregulated | Down syndrome cell adhesion molecule homolog (*Mus musculus*) |
| NV2t005876001.1 | downregulated | Plectin (*Homo sapiens*) |
| NV2t023480001.1 | downregulated | Down syndrome cell adhesion molecule homolog (*Gallus gallus*) |
| NV2t025565001.1 | downregulated | Neuronal cell adhesion molecule (*Gallus gallus*) |
| Nv2t017533001.1 | downregulated | Cadherin-related tumor suppressor (*Drosophila melanogaster*) |
| NV2t006698001.1 | downregulated | Cadherin-related tumor suppressor (*Drosophila melanogaster*) |
